# Therapeutic Potential of PRMT1 as a Critical Survival Dependency Target in Multiple Myeloma

**DOI:** 10.1101/2025.01.29.635603

**Authors:** Tabish Hussain, Sharad Awasthi, Farid Shahid, S. Stephen Yi, Nidhi Sahni, C Marcelo Aldaz

## Abstract

Multiple myeloma (MM) is a neoplasm of antibody-producing plasma cells and is the second most prevalent hematological malignancy worldwide. Development of drug resistance and disease relapse significantly impede the success of MM treatment, highlighting the critical need to discover novel therapeutic targets. In a custom CRISPR/Cas9 screen targeting 197 DNA damage response-related genes, Protein Arginine N-Methyltransferase 1 (PRMT1) emerged as a top hit, revealing it as a potential therapeutic vulnerability and survival dependency in MM cells. PRMT1, a major Type I PRMT enzyme, catalyzes the asymmetric transfer of methyl groups to arginine residues, influencing gene transcription and protein function through post-translational modification. Dysregulation or overexpression of PRMT1 has been observed in various malignancies including MM and is linked to chemoresistance. Treatment with the Type I PRMT inhibitor GSK3368715 resulted in a dose-dependent reduction in cell survival across a panel of MM cell lines. This was accompanied by reduced levels of asymmetric dimethylation of arginine (ADMA) and increased arginine monomethylation (MMA) in MM cells. Cell cycle analysis revealed an accumulation of cells in the G0/G1 phase and a reduction in the S phase upon GSK3368715 treatment. Additionally, PRMT1 inhibition led to a significant downregulation of genes involved in cell proliferation, DNA replication, and DNA damage response (DDR), likely inducing genomic instability and impairing tumor growth. This was supported by Reverse Phase Protein Array (RPPA) analyses, which revealed a significant reduction in levels of proteins associated with cell cycle regulation and DDR pathways. Overall, our findings indicate that MM cells critically depend on PRMT1 for survival, highlighting the therapeutic potential of PRMT1 inhibition in treating MM.

## Introduction

Multiple myeloma, characterized by the malignant proliferation of plasma cells in the bone marrow, accounts for approximately 10% of hematologic cancers and poses a significant therapeutic challenge due to its genetic and phenotypic heterogeneity ^1^. Over the past few decades, use of proteasome inhibitors, immunomodulatory agents, and monoclonal antibodies has significantly increased the five-year survival rate of MM patients to over 50% ^2^. However, despite significant advancements in novel therapies and treatment options, MM remains largely incurable, with almost all patients eventually relapsing or developing resistance to treatment over time ^3^. The persistent challenge of overcoming therapeutic resistance underscores the critical need for novel and more effective treatment strategies.

MM is characterized by profound genomic instability throughout its pathogenesis and progression, making DDR pathways crucial for tumor cell survival. Emerging studies highlight the pivotal role of DDR in driving genomic alterations, sustaining tumor progression, and contributing to therapy resistance in MM ^4–6^. In addition, epigenetic modifications, such as DNA methylation, histone modifications, and RNA-based mechanisms are increasingly recognized as key contributors to the pathogenesis of MM ^7^. Targeting these mechanisms, therefore, presents a promising strategy to address the therapeutic challenges posed by MM.

Here, we employed a custom CRISPR/Cas9 screen targeting 197 DDR-related genes to uncover potential therapeutic vulnerabilities that MM cells rely on for survival. PRMT1 was identified as a top hit, highlighting its critical role in maintaining MM cell viability. As a predominant Type I protein arginine methyltransferase, PRMT1 catalyzes ∼85% of asymmetric dimethylation of arginine residues on histone and non-histone proteins ^8^. Through this post-translational modification, it regulates gene transcription, protein function, and cellular signaling pathways— key processes often dysregulated in cancer ^8–10^. PRMT1 also modulates genome integrity by methylating critical DNA repair proteins, and its dysregulation has been linked to poor prognosis and therapy resistance in both solid tumors and hematological malignancies ^11–13^.

Our study demonstrates that pharmacological inhibition of PRMT1 using the selective Type I PRMT inhibitor GSK3368715 significantly reduced cell survival and proliferation across multiple MM cell lines. Mechanistic analysis revealed that PRMT1 inhibition resulted in cell cycle arrest, reduced levels of ADMA, and downregulation of genes involved in cell proliferation, DNA replication, and DDR. These findings underscore the reliance of MM cells on PRMT1 for survival and suggest that targeting PRMT1 could represent a novel therapeutic strategy in MM. Thus, it provides a strong rationale for the further development of PRMT1 inhibitors in MM treatment and highlights their potential to overcome drug resistance and improve patient outcomes.

## Material and methods

### Cell culture

MM cell lines JJN3, NCI-H929 (RRID:CVCL_1600), KMS11 (RRID:CVCL_2989), and MM1 were cultured in RPMI-1640 medium (ThermoFisher Scientific) supplemented with 10% FBS (Corning), 2 mM L-glutamine, 1 mM sodium pyruvate, and 50 μM β-mercaptoethanol (Sigma). MOPL8, RPMI-8226 (RRID:CVCL_0014), and U266 (RRID:CVCL_0566) cell lines were cultured in RPMI-1640 medium with 10% FBS, 2 mM L-glutamine, and 1 mM sodium pyruvate. All cell lines were maintained at 37°C in a humidified incubator with 5% CO₂.

### CRISPR/Cas9 library screening

A pooled CRISPR/Cas9 sgRNA library targeting 197 genes involved in DDR pathways was a gift from Dr. Simona Colla at MDACC ^14^. It was designed by Cellecta, containing 10 sgRNAs per gene (Supplementary Table S1), and cloned into the pLentiGuide-Puro lentiviral vector. JJN3 cells were transduced with lentivirus generated using the LentiCas9-Blast vector (#52962, Addgene) and selected with 5 μg/mL blasticidin to establish stable JJN3-Cas9 cells. Ten million JJN3-Cas9 cells were then transduced with the pooled sgRNA library at a multiplicity of infection (MOI) < 0.3, ensuring a coverage of 1000x. A total of 5 × 10⁶ cells were collected 48 hours post-transduction as a reference sample (D0). After selection with 1 μg/mL puromycin, cells were harvested again after 21 days (D21). All samples were prepared in triplicate for both D0 and D21. Genomic DNA was isolated using the DNeasy Blood & Tissue Kit (Qiagen) following the manufacturer’s protocol. The sgRNA sequences were amplified using flanking primers with Q5 Hot Start HiFi PCR Master Mix (NEB) and prepared for next-generation sequencing through two rounds of PCR, as described previously ^15^. In the first round, 5 μg of DNA template was used to amplify the sgRNA cassette. In the second round, Illumina sequencing adapters and barcodes were attached through 12 cycles of PCR. The amplified PCR products were purified using the QIAquick PCR Purification Kit, and the expected ∼370 bp band was excised and gel-purified. Quantification of the purified PCR product was performed using a Qubit Fluorometer and Bioanalyzer 2100 (Agilent). The pooled Illumina library was then subjected to NextSeq550 high-output sequencing with >1000x coverage per sample.

### CRISPR screening data analysis

The sgRNA sequences were pulled out from the paired end sequence files using the FASTX-Toolkit (http://hannonlab.cshl.edu/fastx_toolkit/index.html). For each sgRNA in the library, the count of mapped reads was calculated for each time point, D0 or D21. A maximum-likelihood estimation (MLE) of the gene essentiality score for each gene was generated using the Model-based Analysis of Genome-wide CRISPR/Cas9 Knockout (MAGeCK) algorithm ^16^. To identify high-confidence hits, we employed a stringent false discovery rate (FDR) threshold. It revealed a set of significantly depleted genes, which we defined as the core target genes.

### Long term cell survival assay

The Type I PRMT inhibitor GSK3368715, with a purity of 99.96%, was procured from MedChemExpress (HY-128717). JJN3 and NCI-H929 cells (2,000 cells per well) were seeded in a 96-well plate and treated with either DMSO or various concentrations of GSK3368715: 0.004 µM, 0.01 µM, 0.04 µM, 0.12 µM, 0.37 µM, 1.1 µM, 3.3 µM, 10 µM, or 20 µM for 14 days. After the treatment period, cell viability was assessed using the CellTiter-Glo kit (Promega) according to the manufacturer’s instructions. Results were quantified as a percentage of viability relative to the DMSO-treated control and used to calculate gIC50.

### Immunoblot analysis

JJN3 and NCI-H929 cells were treated with either DMSO, 0.1 µM, 1 µM, or 10 µM GSK3368715 for 48 hours. Cells were lysed using RIPA buffer supplemented with protease and phosphatase inhibitors (Roche) and 0.25 mM phenylmethanesulfonyl fluoride (PMSF) (Sigma). Protein concentrations were quantified using the BCA assay (Promega), and equal amounts of protein were loaded onto a 12% SDS-PAGE gel. After electrophoresis, proteins were transferred to a PVDF membrane. The following antibodies were used for detection: PRMT1 (Cell Signaling Technology, CST #2449, RRID:AB_2237696, 1:1000), Mono-Methyl Arginine (MMA-RGG) (CST #8711, RRID:AB_10896849, 1:1000), Asymmetric Di-Methyl Arginine Motif (ADMA) (CST #13522, RRID:AB_2665370, 1:1000), and HSP90 (CST #4874, RRID:AB_2121214, 1:1000).

### Cell cycle analysis

Cell cycle analysis was performed as previously described ^17^. Briefly, JJN3 and NCI-H929 cells were treated with either DMSO, 1 µM, or 10 µM GSK3368715 for 72 hours and fixed in 70% ethanol. After fixation, cells were washed twice with PBS and stained with a buffer containing 50 µg/mL propidium iodide (Sigma) and 0.5 mg/mL RNaseA (Sigma). To prevent cell clumping, samples were passed through a syringe before data acquisition on a flow cytometer using 488 nm excitation. Cell cycle phase distribution was analyzed using FlowJo 10 software (RRID:SCR_008520).

### Apoptosis assay

Cell death by apoptosis was measured by Annexin V-PI staining as previously described ^17^. Briefly, JJN3 and NCI-H929 cells were treated with either DMSO, 0.1 µM, 1 µM, or 10 µM GSK3368715 for 14 days. Cells were washed with PBS and incubated with Annexin V, Alexa Fluor™ 647 conjugate (1:100 dilution, ThermoFisher Scientific) and PI (1 μg, Sigma) prepared in incubation buffer (10 mM Hepes/NaOH, pH 7.4, 140 mM NaCl, 5 mM CaCl_2_) for 15 min and analyzed by flow cytometry at 650 nm excitation.

### Quantitative RT-PCR

Quantitative RT-PCR (qRT-PCR) was performed as previously described ^18^. Briefly, total RNA extracted from JJN3 and NCI-H929 cells treated with either DMSO, 1 µM, and 10 µM GSK3368715 for 72 hours. cDNA was synthesized using High-Capacity cDNA Reverse Transcription Kit (Applied Biosystems) following manufacturer’s instructions. The relative expression level for specific genes was determined in triplicate by qRT-PCR using the SYBR Green-based method. After normalization to 18s RNA expression, the average fold change was calculated using the 2^-(ΔΔCt) method described elsewhere ^19^. Sequence of the primers used for qRT-PCR is given in Supplementary Table S2.

### Reverse-phase protein array (RPPA)

JJN3 cells were treated with either DMSO, 1 µM, or 10 µM GSK3368715 for 72 hours. Cell lysates were prepared using RIPA buffer containing protease and phosphatase inhibitors (Roche), then mixed with 4x SDS buffer. Samples were boiled for 5 minutes, snapped frozen, and processed at the RPPA Core Facility at MD Anderson Cancer Center, as previously described ^20^. Each sample was analyzed in triplicate. Protein expression values from RPPA analysis were compared between GSK3368715 treated and control groups using a t-test, followed by p-value adjustment for multiple testing to calculate FDR. Statistically significant proteins (p < 0.05) were selected, and Log2 FC was calculated for further analysis using Ingenuity Pathway Analysis (IPA) (RRID:SCR_008653).

### Statistical analysis of quantitative RT-PCR, cell cycle, and apoptosis assays

Statistical analyses were performed using GraphPad Prism 10 (RRID:SCR_002798), employing one-way ANOVA with Tukey’s and Dunnett’s post-hoc tests for multiple comparisons. Each experiment included at least three replicates. Data are presented as mean values ±SEM. The number of replicates for each experiment is indicated in the figure legends and represented by data points in the figures.

## Results

### *In vitro* CRISPR screen identified *PRMT1* as a key genetic vulnerability in MM

To uncover novel survival dependencies, we performed a CRISPR-Cas9 loss-of-function screen in the JJN3 MM cell line, aiming to identify genes whose disruption is lethal to these cells. Our overall strategy involved transducing JJN3 cells with the pooled lentiviral library and sequencing the sgRNAs from viable cells at two time points: after transduction (D0) and after 21 days of puromycin selection (D21). This approach was designed to identify survival dependency targets, with the expectation that sgRNAs targeting essential genes for MM cell survival would be depleted by D21 compared to D0. We used a custom CRISPR library of ∼2,000 sgRNAs targeting 197 DDR-related genes (about 10 sgRNAs per gene). To ensure thorough coverage, we transduced ∼ 10×10^6^ cells to maintain 1000× coverage for each sgRNA throughout the experiment. Following transduction, JJN3 cells were selected with puromycin for stable viral integration and cultured for 21 days to eliminate cells with essential genetic variation for MM survival. Deep sequencing of sgRNAs from DNA isolated from viable cells at D0 and D21 was then performed to identify sgRNAs depleted by D21 (Figure 1a). Analysis using the MAGeCK algorithm identified a set of 10 fitness genes (FDR < 0.05) crucial for the viability of JJN3 cells (Figure 1b). PRMT1 was identified as a leading candidate with a remarkably low Robust Rank Aggregation (RRA) score of 0.0000336, showing statistical significance (p = 0.0003, FDR = 0.01). Due to its established role in cancer, this finding prompted further investigation into PRMT1 as a potential therapeutic vulnerability and survival dependency target in MM cells.

**Figure 1.**
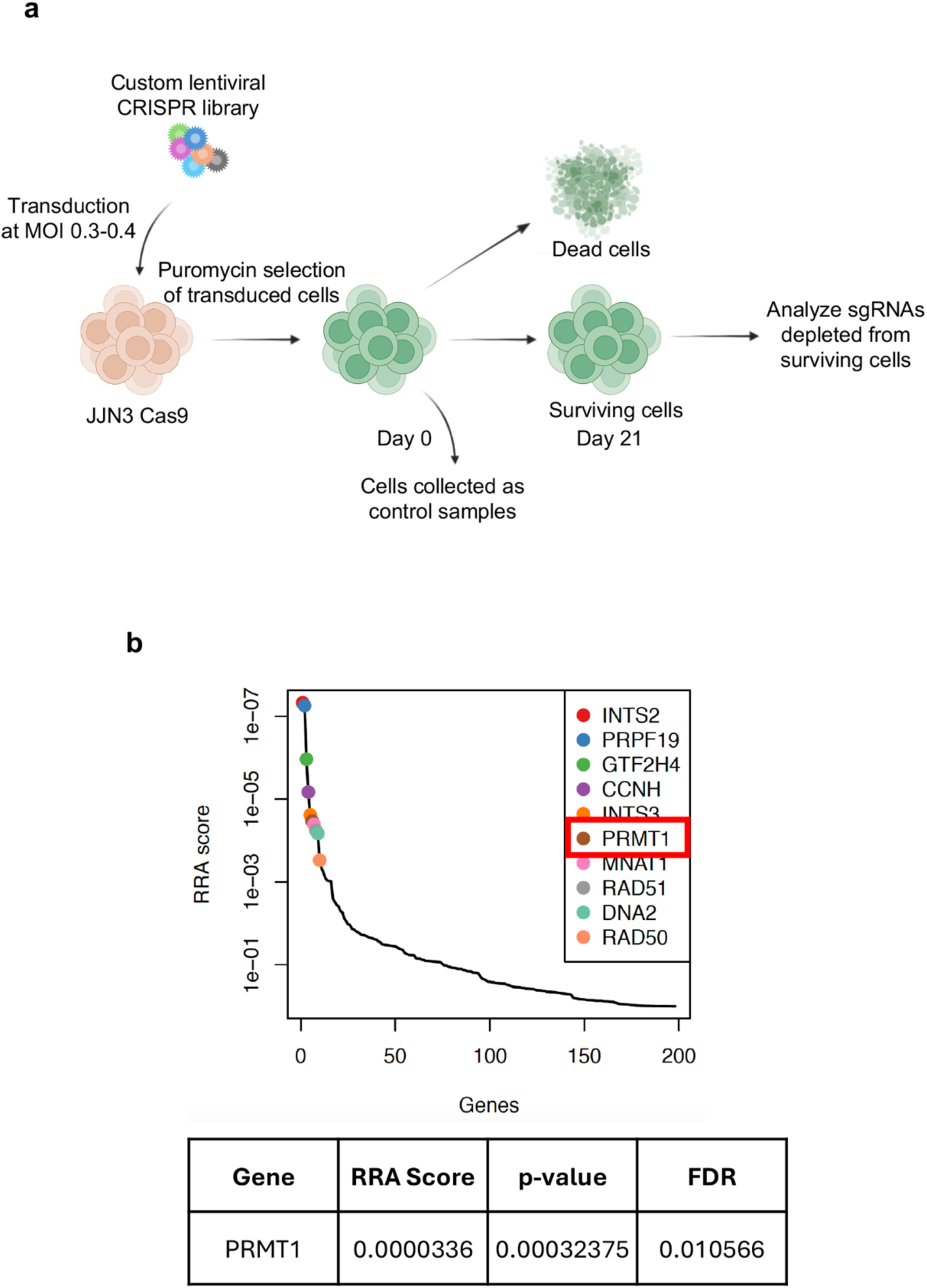
CRISPR screening identified *PRMT1* as key survival dependency target in MM. **(a)** Schematic of the CRISPR/Cas9 negative knockout screening workflow. A custom CRISPR/Cas9 knockout library (∼2000 sgRNAs targeting 197 DDR-related genes) was packaged into lentiviral particles and transduced into Cas9-expressing JJN3 cells (JJN3-Cas9) at a low multiplicity of infection (MOI: 0.3–0.4). Transduced cells were selected with puromycin and cultured for 21 days to eliminate cells with essential genetic variations critical for MM survival. **(b)** sgRNA abundance was quantified via high-throughput sequencing and analyzed using the MAGeCK algorithm. *PRMT1* emerged as a leading candidate with a low RRA score (0.0000336) and strong significance (p = 0.0003, FDR = 0.01).

### PRMT1 inhibition reduces MM cell viability and triggers cell cycle progression defects

To assess the effect of PRMT1 inhibition on MM cells growth and viability, we utilized GSK3368715, a potent Type I PRMT inhibitor (hereafter referred to as PRMTi) ^21^. A 14-day cell viability assay was conducted across seven MM cell lines, generating dose-response curves relative to DMSO-treated controls, and determining the growth half-maximal inhibitory concentration (gIC50) (Figure 2a). The analysis revealed varying sensitivities to PRMTi across MM lines, with gIC50 values ranging from as low as 0.021 μM for the most sensitive cell line, NCI-H929, to 14.405 μM for the least sensitive cell line, MM1 (Figure 2a). Based on these results, we selected JJN3 and NCI-H929 for further investigation. PRMT1 is the key methyltransferase responsible for the generation of ADMA. Consequently, effective PRMT1 inhibition leads to decreased global levels of ADMA and an increase in global MMA ^22^. Using methylarginine-specific antibodies, we observed a significant dose dependent reduction in ADMA levels and a concomitant increase in MMA in JJN3 and NCI-H929 cells with PRMT1 depletion, compared to DMSO-treated control, indicating efficient PRMT1 inhibition (Figure 2b, Supplementary Figure S1). Western blot analysis revealed a distinct banding pattern, highlighting the broad spectrum of proteins modified post-transcriptionally by PRMT1 (Figure 2b).

**Figure 2.**
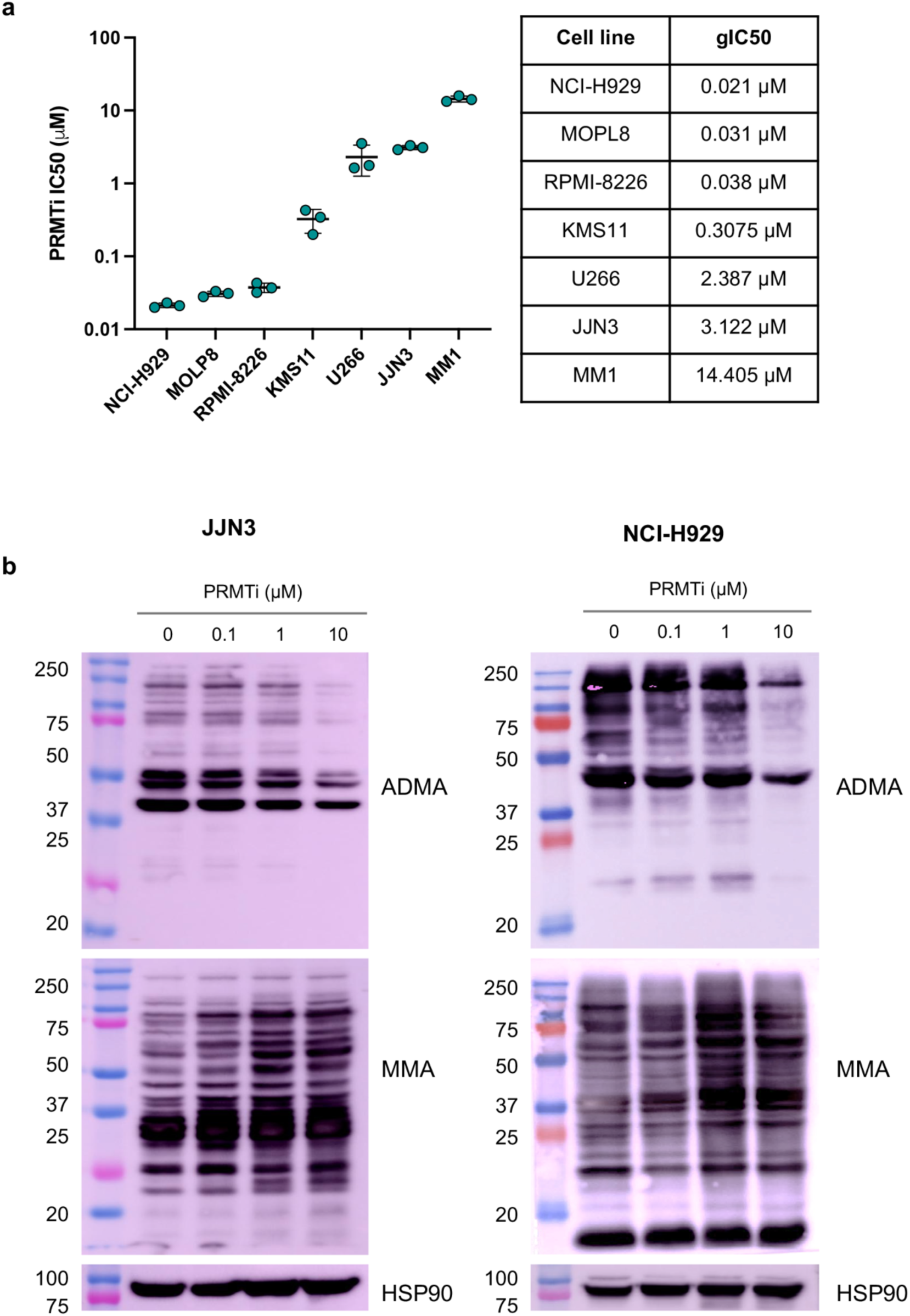
PRMT1 inhibition impairs MM cell viability, reduces ADMA, and increases MMA. **(a)** PRMTi gIC50 values for a panel of MM cell lines determined using dose-response curves from cell survival assays. Each data point represents the gIC50 calculated from individual replicates for each cell line. Data are presented as mean ± SEM from three replicates per cell line. (**b**) Western blot analysis showing the impact of PRMTi on total asymmetric arginine dimethylation (ADMA) and arginine monomethylation (MMA) in JJN3 and NCI-H929 MM cell lines treated with DMSO or different concentrations of PRMTi for 48 hours.

To evaluate cell proliferation and clonogenic survival of MM cells, we performed a limiting dilution assay with various cell densities across different PRMTi treatment concentrations. This assay allowed us to quantify both the proliferation and survival capacity of MM cells under conditions of limited cell density. The results showed a dose- and cell density-dependent reduction in cell proliferation and clonogenic potential following PRMTi treatment (Figure 3a). We further evaluated the cell cycle profile of PRMTi-treated MM cells and observed a significant accumulation of cells in the G0/G1 phase, with a simultaneous reduction in the S phase at 72 hours post-treatment in both MM lines (Figure 3b and c). This effect was statistically significant at 1 µM PRMTi and intensified further at higher PRMTi concentration (Figure 3b and c). We also performed Annexin V-PI staining which revealed that PRMT1 inhibition did not increased apoptotic cell death (Supplementary Figure S2). These findings suggest that while PRMT1 inhibition impairs cell growth and disrupts cell cycle progression, it does not activate the apoptotic pathway in these MM cells.

**Figure 3.**
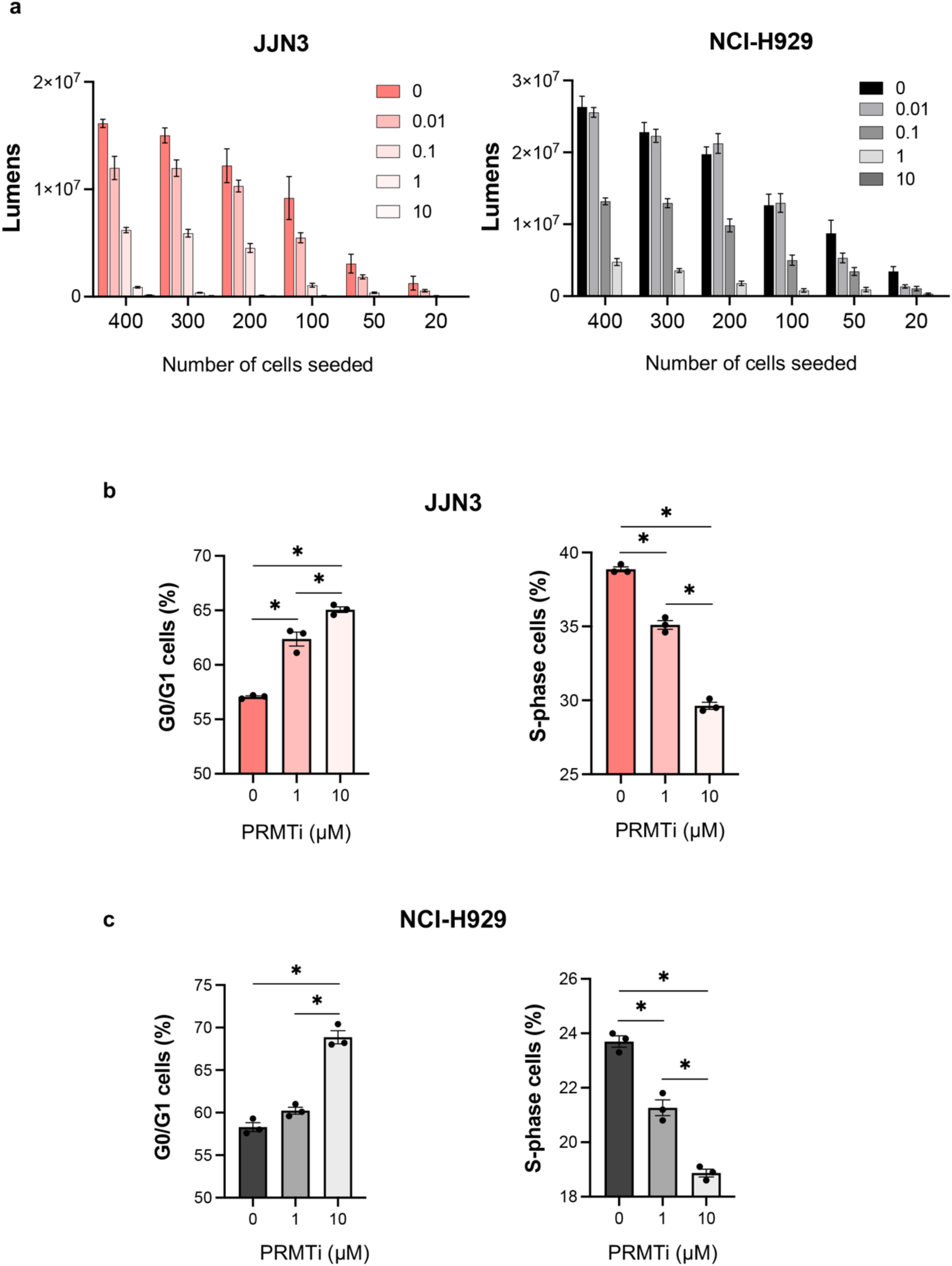
PRMT1 inhibition reduces clonogenic potential and triggers cell cycle defects in MM cells. (**a**) Limiting dilution assay showing a dose- and cell density-dependent decrease in cell proliferation and clonogenic potential in JJN3 and NCI-H929 cells following PRMTi treatment. Bars represent the mean ± SEM of 10 replicates for each cell number and dose concentration. (**b**) Bar graph showing the percentage of cells arrested in G0/G1 and S phases 72 hours post-PRMTi treatment in JJN3 and NCI-H929 cells. Data are presented as the mean ± SEM from three replicates (*= p < 0.05, One-way ANOVA with Tukey’s multiple comparison test).

### PRMT1 regulates key genes involved in cell cycle regulation, DNA replication, and repair mechanisms

Given that PRMTi treatment reduced MM cell survival and lead to disruption in cell cycle progression, we sought to further investigate the molecular effects of PRMT1 inhibition on critical cellular processes. To this end, we performed qRT-PCR analyses targeting key genes associated with DNA replication, cell cycle regulation, and DDR. PRMT1 inhibition led to significant downregulation of essential genes involved in DNA replication. Specifically, *CDC45*, which plays a crucial role in the initiation of DNA replication ^23^, and *PRIM1*, required for the synthesis of RNA primers ^24^, were markedly reduced in expression (Figure 4a). The expression of key regulators of the cell cycle was also notably affected by PRMT1 inhibition. *CLSPN*, which is integral to the checkpoint response during DNA replication ^25^, and *TOPBP1*, which facilitates the ATR-mediated checkpoint signaling ^26^, were significantly downregulated (Figure 4b). This indicates that PRMT1 inhibition disrupts the DNA replication machinery and is crucial for maintaining proper cell cycle progression and checkpoint control, likely leading to cell cycle arrest and reduced MM cell proliferation. Genes critical for DDR also exhibited reduced expression following PRMT1 inhibition. *BRCA1* and *BRCA2*, which are critical for homologous recombination repair of double-strand DNA breaks ^27^, along with *FANCD2*, essential for DNA interstrand cross-link repair ^28^, were significantly reduced (Figure 4c). Additionally, *FEN1*, involved in DNA replication and repair ^29^, *ERCC4*, a key component of nucleotide excision repair ^30^, and *RAD51*, which plays a crucial role in homologous recombination ^31^, also showed reduced expression levels (Figure 4c). Collectively, these results highlight the broad impact of PRMT1 inhibition on essential cellular functions. The downregulation of key genes within these pathways provides a mechanistic explanation for the observed deficits in cell viability and proliferation in PRMTi treated MM cells.

**Figure 4.**
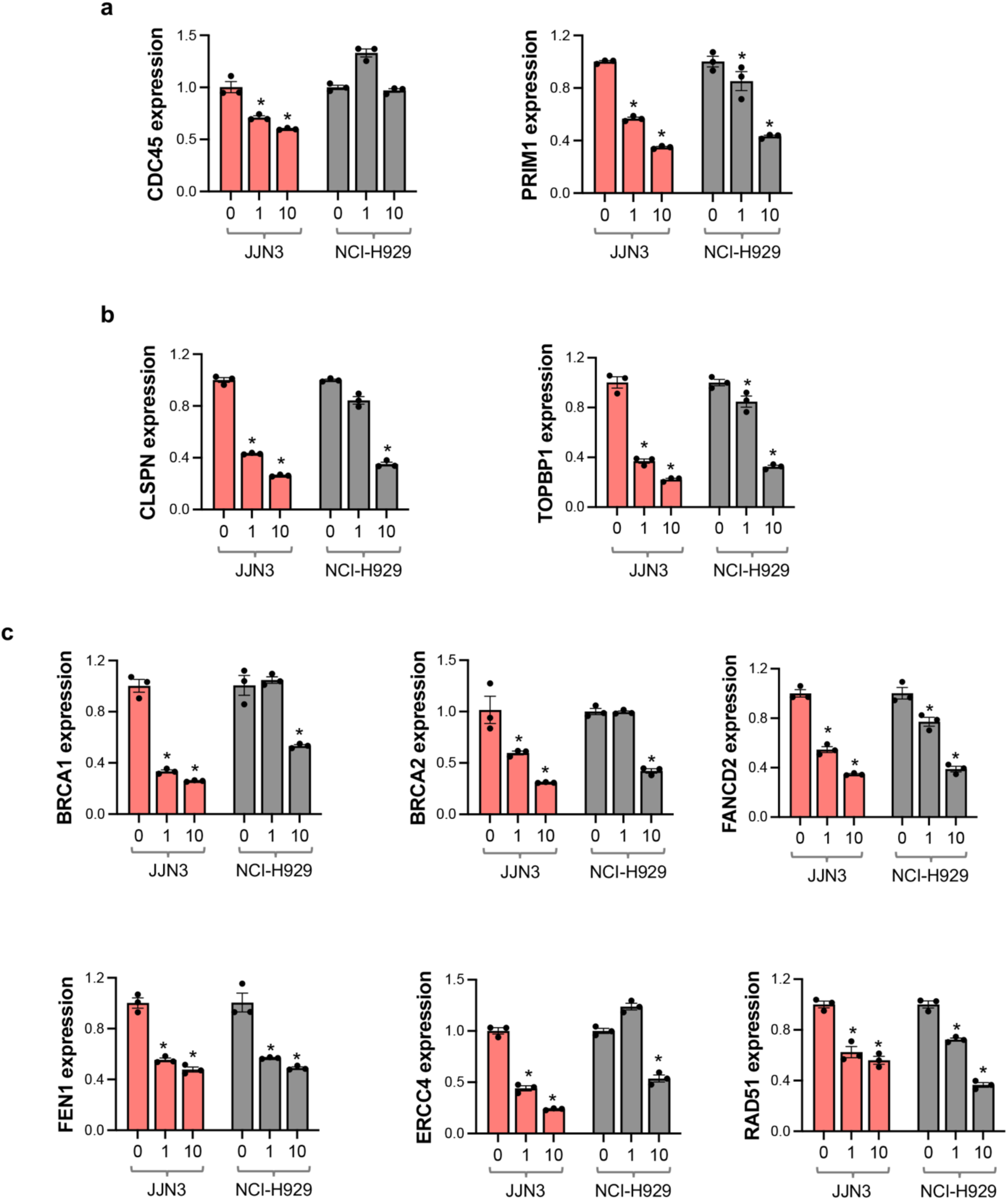
PRMT1 regulates genes critical for cell cycle regulation, DNA replication, and repair. (**a-c**) Bar graphs showing the effect of PRMTi treatment on mRNA expression levels, measured by qRT-PCR, on DNA replication genes *CDC45* and *PRIM1* (**a**), cell cycle regulators *CLSPN* and *TOPBP1* (**b**), and DNA damage repair genes *BRCA1*, *BRCA2*, *FANCD2*, *FEN1*, *ERCC4*, and *RAD51* (**c**) in JJN3 and NCI-H929 cells. Data are presented as mean ± SEM from three independent replicates (*= p < 0.05, One-way ANOVA with Dunnett’s multiple comparison test).

### Protein level dysregulation of cell proliferation, cell cycle, and DDR pathways after PRMT1 inhibition

To further explore the impact of PRMT1 inhibition on critical molecular pathways, we performed Reverse Phase Protein Array (RPPA) analyses on JJN3 cells exposed to 1 µM and 10 µM PRMTi concentrations. This allowed a comprehensive profiling of ∼500 proteins and phosphoproteins involved in critical cellular processes, including cell cycle regulation, proliferation, signal transduction, apoptosis, metastasis, DNA replication, DDR, and metabolism. A strong positive correlation was observed between the differential expression profiles of proteins and phosphoproteins (DEPs) (Pearson correlation coefficient = 0.7151), indicating that they were consistently regulated in the same direction, either upregulated or downregulated, across the two doses (Figure 5a). At 1 µM PRMTi, 109 proteins were differentially regulated (p < 0.05), while at 10 µM PRMTi, the number increased to 316 proteins (p < 0.05) (Figures 5b and c, Supplementary File S1). A greater number of DEPs at the higher PRMTi dose indicates an impact on a broader range of proteins, suggesting a more significant overall impact. Unsupervised hierarchical clustering of DEPs effectively differentiated between the PRMT1i 1 µM vs Ctrl (Figure 5b) and PRMTi 10 µM vs Ctrl (Figure 5c) treatment groups.

**Figure 5.**
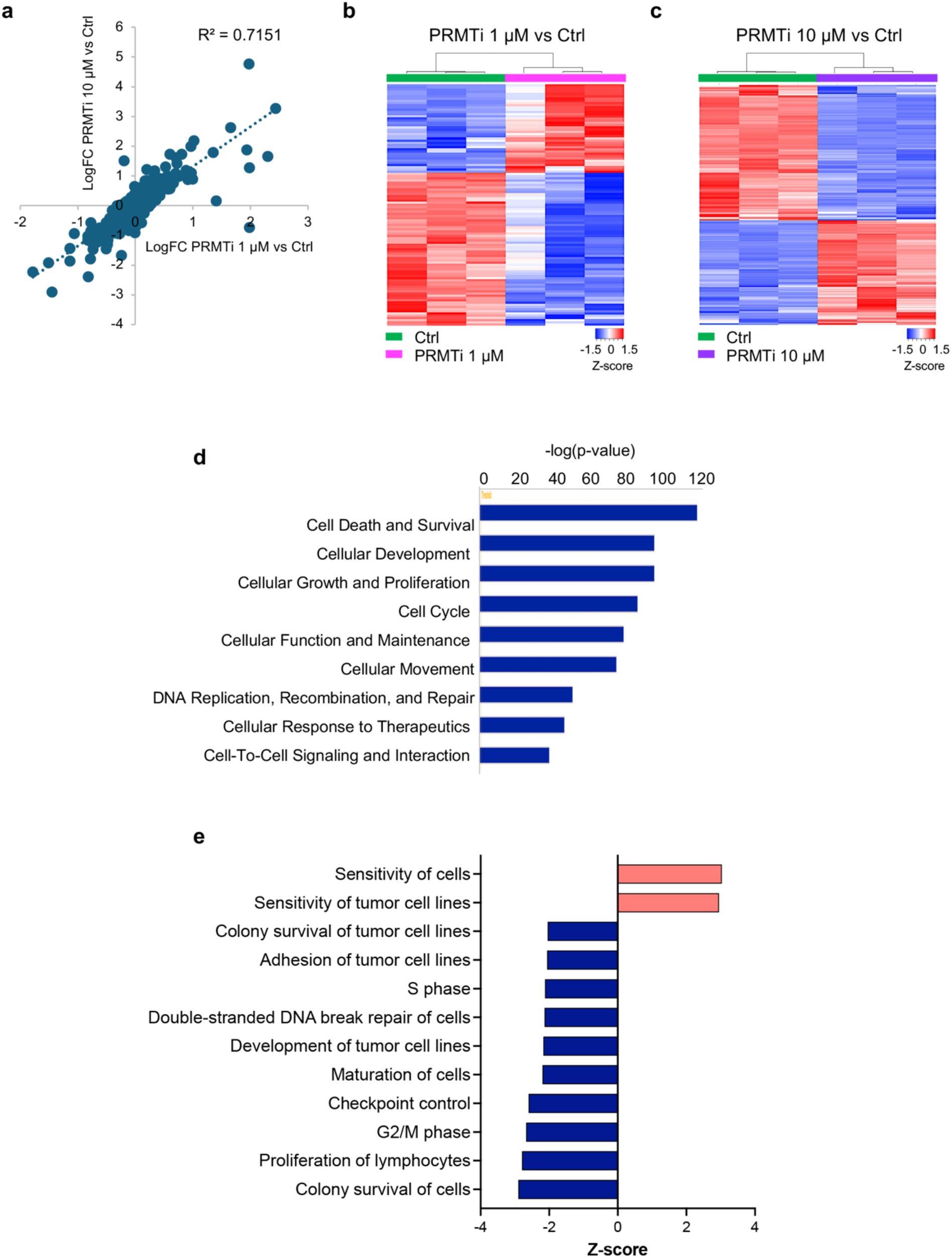
PRMT1 inhibition disrupts protein expression, impacting critical cellular functions. (**a**) Scatter plot showing a strong positive correlation between differential protein expression profiles as per RPPA analyses at 1 µM and 10 µM PRMTi concentrations (Pearson correlation coefficient = 0.7151). (**b-c**) Unsupervised hierarchical clustering of RPPA profiles showing distinct separation between control and PRMTi-treated JJN3 MM cells (n = 3 replicates per group). Heatmaps displaying profiles of 109 differentially expressed proteins (FDR < 0.05) comparing PRMTi 1 µM vs. ctrl (**b**) and 316 differentially expressed proteins (FDR < 0.05) comparing PRMTi 10 µM vs. ctrl (**c**). Proteins with increased or decreased expression levels are represented in red and blue, respectively. (**d**) Bar graph showing IPA-based enrichment of key molecular and cellular functions in PRMTi-treated JJN3 cells. (**e**) Bar graph showing negatively enriched functions, including Colony survival of cells, Proliferation of lymphocytes, DNA damage repair, cell cycle progression, and checkpoint control. Positively enriched functions indicate increased sensitivity to therapeutics in PRMTi-treated JJN3 cells.

We analyzed the RPPA expression data of 316 DEPs in the PRMTi 10 µM vs Ctrl group and calculated Log2 fold change, p-value, and FDR. Among these DEPs, 138 were upregulated, while 178 were downregulated relative to the Ctrl group (p < 0.05, FDR < 0.05) (Supplementary File S1). This dataset was utilized for Ingenuity Pathway Analysis (IPA), which revealed significant enrichment in several key molecular and cellular functions, including ‘Cell death and survival’, ‘Cellular growth and proliferation’, ‘Cell cycle’, ‘DNA replication and damage repair’ and ‘cellular response to therapeutics’, associated with the reversal of tumorigenesis (Figure 5d). Within these categories the functions that were negatively enriched included Colony survival of cells, Proliferation of lymphocytes, Development of tumor cell lines, and Adhesion of tumor cell lines (Z score range: −2.9 – −2.04, p value range: 3.9E-55 – 1.93E-30, Figure 5e). Additionally, there was a notable negative impact on DNA damage repair, cell cycle progression (S phase and G2/M phase), and checkpoint control (Z score range: −2.67 – −2.12, p value range: 2.48E-45 – 1.05E-23, Figure 5e). In contrast, functions with a positive Z-score were associated with increased sensitivity in PRMTi-treated cells, highlighting an enhanced cellular response to therapeutics in JJN3 cells (Z score = 3.03, p value = 1.33E-47, Figure 5e).

Comparative analysis of DEPs between 1 µM and 10 µM PRMTi treatment groups identified 102 overlapping proteins (Figure 6a, Supplementary File S1). These common proteins exhibited a highly consistent expression pattern, with a strong positive correlation (Pearson correlation coefficient = 0.8638) (Figure 6b). Functional annotation revealed that these shared DEPs were primarily associated with DDR, cell death and apoptosis, cell cycle regulation, metabolism, transcription regulation, tumor metastasis, DNA replication, and epigenetic regulation. A significant proportion of these DEPs also belonged to various cell signaling pathways (Supplementary Figure S3). Most differentially expressed DDR proteins showed reduced expression with 10 µM PRMTi treatment (Figure 6c). Similarly, DEPs associated with cell cycle regulation, DNA replication, transcription regulation, and cell signaling were consistently downregulated at this dose (Figure 6c). These findings align with cellular assays and qRT-PCR data, which revealed significant suppression of DNA replication, cell cycle regulation, and DDR pathways. This data further highlighted that PRMT1 inhibition disrupts these processes, resulting in reduced proliferation and genomic instability in multiple myeloma cells. Interestingly, proteins involved in metabolic regulation showed increased expression at the higher PRMTi dose, suggesting potential compensatory metabolic reprogramming to support cell survival under stress induced by PRMT1 inhibition (Figure 6c).

**Figure 6.**
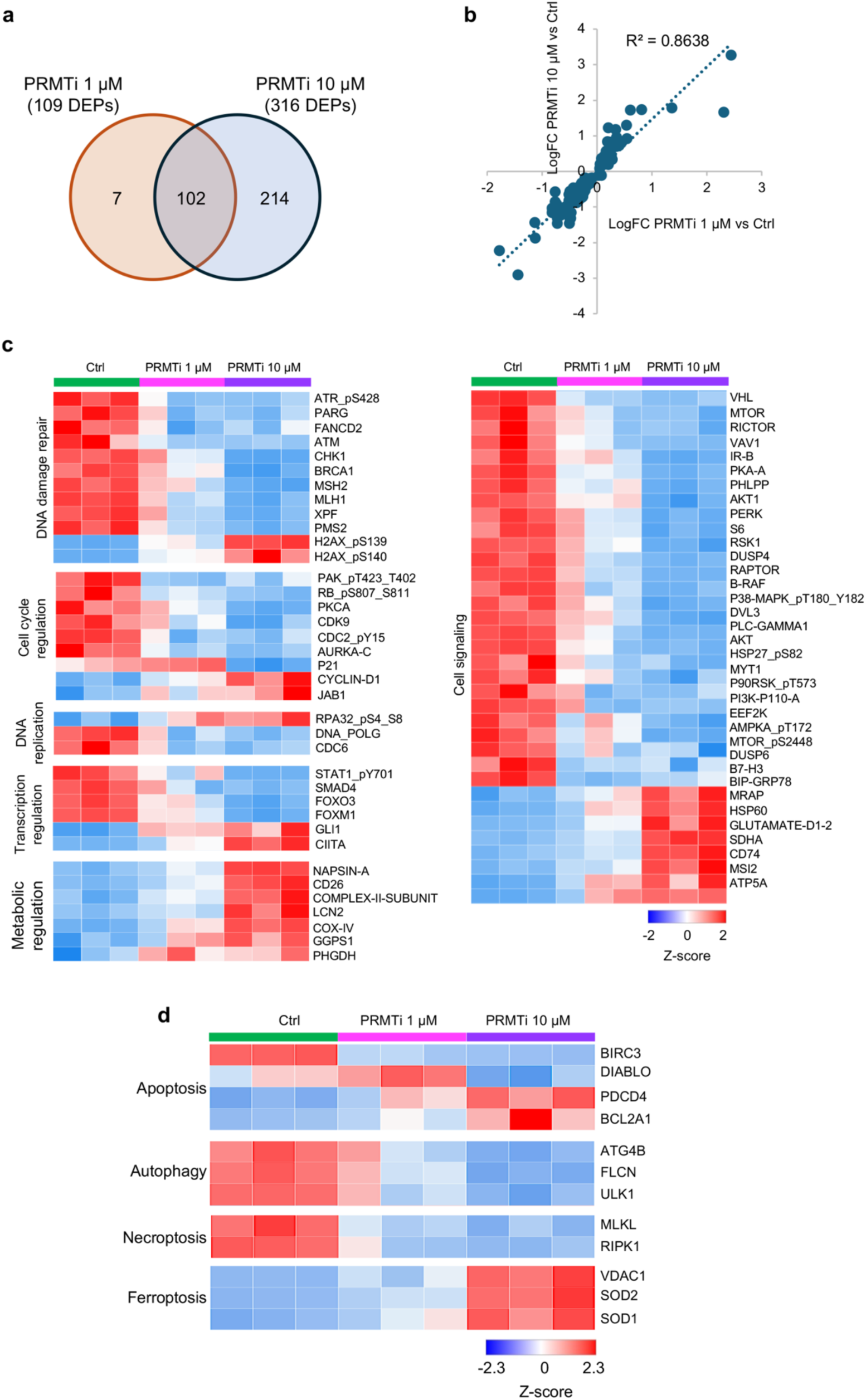
PRMT1 inhibition downregulates key cellular pathways and highlights ferroptosis as a potential cell death mechanism. **(a)** Venn diagram showing the 102 shared differentially expressed proteins (DEPs) between PRMTi 1 µM vs. ctrl and PRMTi 10 µM vs. ctrl groups. **(b)** Scatter plot illustrating a strong positive correlation in differential protein expression profiles for the 102 shared proteins between 1 µM and 10 µM PRMTi concentrations (Pearson correlation coefficient = 0.8638). (**c**) The heatmap shows PRMTi concentration-dependent downregulation of proteins involved in DDR, cell cycle regulation, DNA replication, transcription regulation, and cell signaling pathways in JJN3 MM cells. In contrast, metabolism-related proteins were upregulated, suggesting compensatory metabolic reprogramming in response to PRMT1 inhibition-induced stress. (**d**) The heatmap illustrates PRMTi concentration-dependent effects on proteins associated with various cell death processes, including apoptosis, autophagy, necroptosis, and ferroptosis. Upregulation of ferroptosis-related proteins, such as SOD1, SOD2, and VDAC1, suggests ferroptosis as a potential mechanism of cell death induced by PRMT1 inhibition. Proteins with increased or decreased expression levels are shown in red and blue, respectively.

RPPA data also highlighted effects on proteins involved in various cell death processes, including apoptosis, autophagy, necroptosis, and ferroptosis (Figure 6b). In PRMTi-treated MM cells, the pro-apoptotic protein DIABLO was downregulated, while the anti-apoptotic protein BCL2A1 was upregulated, indicating that apoptosis is unlikely the primary mode of cell death. This aligns with our Annexin V-PI staining results, which showed no evidence of apoptotic cell death following PRMT1 inhibition. Proteins associated with autophagy (ATG4B, FLCN, ULK1) and necroptosis (MLKL, RIPK1) were similarly downregulated, further excluding these mechanisms (Figure 6b). Interestingly however, ferroptosis-related proteins, including SOD1, SOD2, and VDAC1, were upregulated, suggesting that ferroptosis may be the potential mechanism of cell death initiated by PRMT1 inhibition (Figure 6b).

## Discussion

By means of applying a CRISPR-Cas9 loss-of-function screen, we identified PRMT1 as a key genetic vulnerability and survival dependency target in MM cells. Identification of PRMT1 as a top hit underscores its essential role in MM cell survival, with its disruption leading to significant fitness defects. Our findings align with the growing recognition of PRMT1 as a promising therapeutic target, offering a novel avenue for intervention in several cancers, including MM ^13,32–35^.

PRMT1 is the most prominently expressed PRMT in MM cell lines and primary malignant plasma cells from MM patients, with significantly elevated levels observed in relapsed/refractory patients and linked to poor prognosis, highlighting its potential role in disease progression and treatment resistance ^34,35^. GSK3368715 is a potent inhibitor of type I protein arginine methyltransferases, primarily targeting PRMT1, the predominant Type I enzyme. This PRMT1i has been tested *in vitro* across various cancers, demonstrating the most pronounced effects in cell lines from two major hematopoietic malignancies such as lymphoma and acute myeloid leukemia (AML) ^21^. Our study provides a detailed analysis of the effects of PRMT1 inhibition using GSK3368715 on MM cells. Treatment with this specific PRMT1i led to a significant reduction in cell viability across various MM cell lines. JJN3 and NCI-H929 cells displayed dose- and cell density-dependent decreases in cell proliferation and clonogenic potential as a consequence of substantial cell cycle arrest. These observations align with previous studies, where genetic disruption and pharmacological inhibition of PRMT1 using MS023, were shown to have similar effects on MM cells ^34,35^.

Consistent with the cellular phenotypes induced by PRMT1 inhibition, analysis by qRT-PCR showed PRMTi mediated downregulation of specific genes involved in cell cycle progression, DNA replication, and damage repair pathways in MM cells. The critical cell cycle regulators, *CLSPN* and *TOPBP1* and DNA replication genes, *CDC45* and *PRIM1*, exhibited a significant loss of expression upon PRMTi treatment in MM cells. Additionally, DDR genes, including *BRCA1*, *BRCA2*, *FANCD2*, *FEN1*, *ERCC4*, and *RAD51*, integral to various DNA repair pathways, were severely impacted. These findings are consistent with earlier studies showing that PRMT1 loss in mouse embryonic fibroblasts resulted in spontaneous DNA damage, delayed cell cycle progression, checkpoint defects, and chromosomal abnormalities ^36^. Similarly, recent research in pancreatic ductal adenocarcinoma demonstrated that PRMT inhibition using GSK3368715 disrupted cell cycle regulation, DNA replication, and damage response pathways, significantly impairing tumorigenesis ^12^. Our RPPA data also corroborated these findings, showing widespread dysregulation of proteins involved in cell cycle regulation, DNA replication, transcription regulation, and DDR pathways. Together, these data highlight the broad impact of PRMT1 inhibition in MM cells, likely driven by its ability to methylate arginine residues on both histone and non-histone proteins, thereby influencing gene expression through chromatin remodeling and regulating essential processes such as transcription, DNA damage response, and cell signaling.

Despite observing reduction in cell viability and proliferation, we did not detect significant changes in apoptosis. Furthermore, RPPA data revealed decreased levels of proteins involved in apoptosis, autophagy, and necroptosis pathways, suggesting that PRMT1 inhibition by GSK3368715 likely affects an alternative mechanism of cell death. Interestingly, RPPA analysis revealed an upregulation of ferroptosis-related proteins, suggesting that ferroptosis is potentially the underlying mechanism of cell death induced by PRMT1 inhibition. This observation aligns with findings from a recent study in AML, where PRMT1 was identified as a critical ferroptosis regulator and treatment with GSK3368715 enhanced ferroptosis sensitivity *in vitro* and *in vivo* ^37^.

In conclusion, our findings establish PRMT1 as a critical therapeutic vulnerability and survival dependency in MM cells. PRMT1 inhibition disrupted several key pathways ultimately inducing cellular stress and impairing the ability of MM cells to recover or proliferate. This highlights the potential of PRMT1 as a valuable target for therapeutic intervention, either alone or in combination with existing therapies. Future studies exploring combinatorial approaches with PRMT1 inhibitors and contemporary therapies could help improve treatment outcomes in MM.

## Acknowledgement

The authors thank Dr. Simona Colla at MD Anderson Cancer Center for providing the Cellecta CRISPR/Cas9 DDR library, which significantly contributed to this study.

## Author Contributions

T.H.- designed and performed the experiments and manuscript writing; S.A.- provided assistance with the CRISPR screen; F.S.- provided assistance with cell cycle and limiting dilution assays; S.S.Y.- conducted bioinformatics data analysis for the CRISPR screen; N.S.- provided intellectual input and support on CRISPR screening and RPPA assay; and C.M.A. conceptualized the study, manuscript writing, and correspondence

---

**Supplementary Figure S1.**
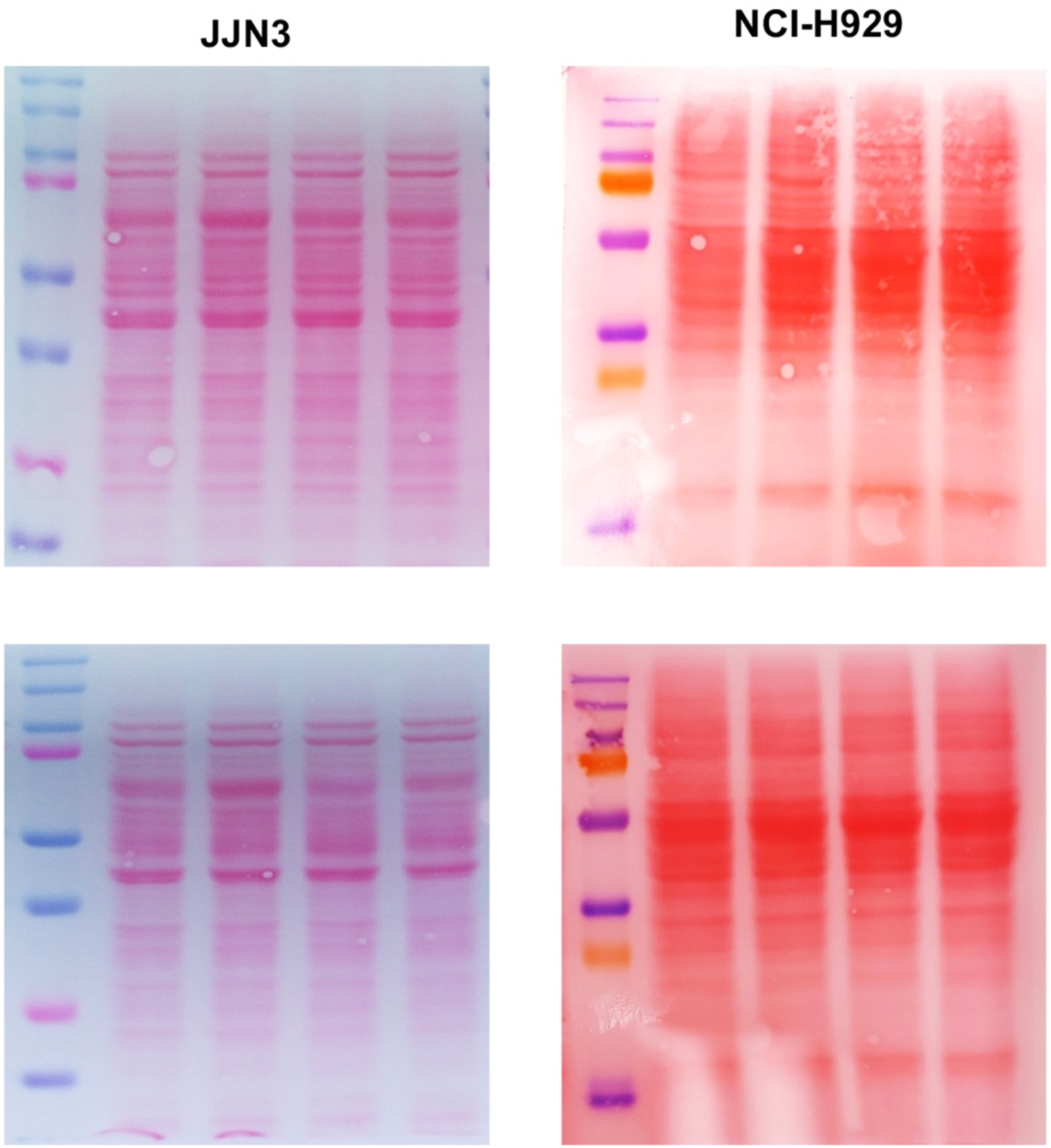
Ponceau staining confirms equal protein loading. Ponceau staining of membranes from ADMA and MMA western blots in JJN3 and NCI-H929 cells, confirming equal protein loading.

**Supplementary Figure S2.**
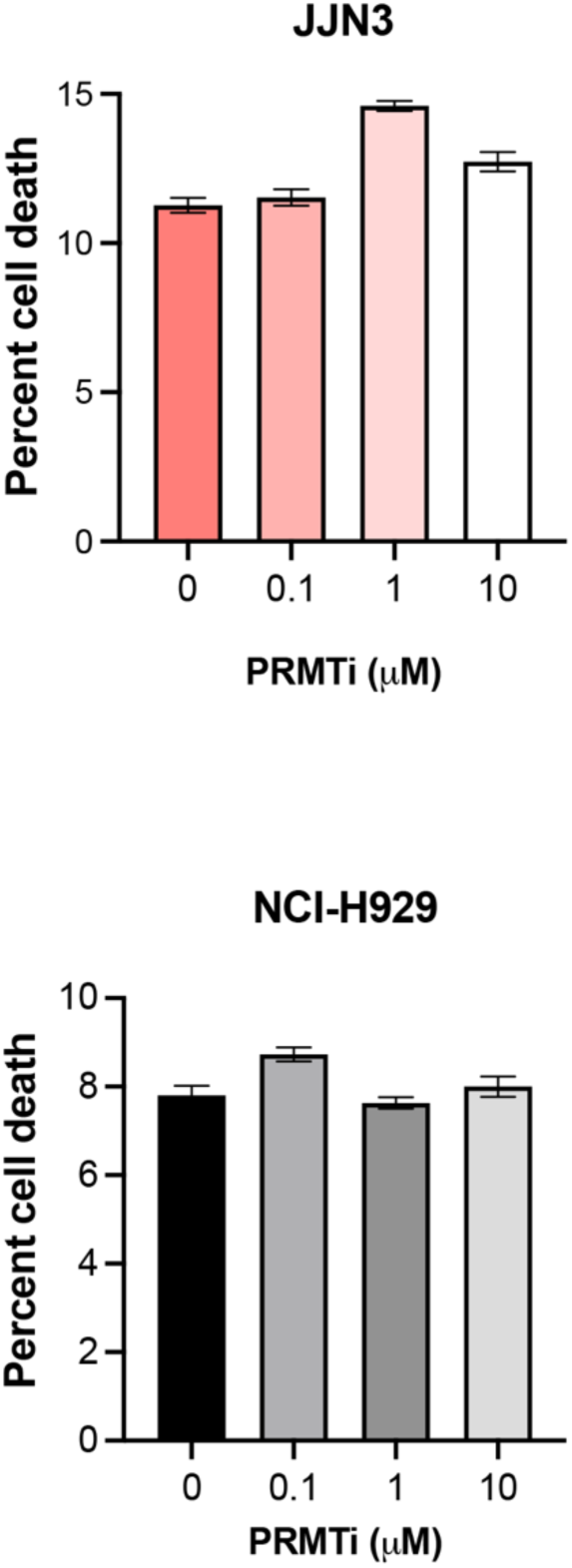
PRMT1 inhibition does not increase apoptotic cell death. Bar graph showing the percentage of apoptosis and cell death measured by Annexin V-PI staining of JJN3 and NCI-H929 MM cells 72 hours post-PRMTi treatment. Data are presented as the mean ± SEM from three replicates.

**Supplementary Figure S3.**
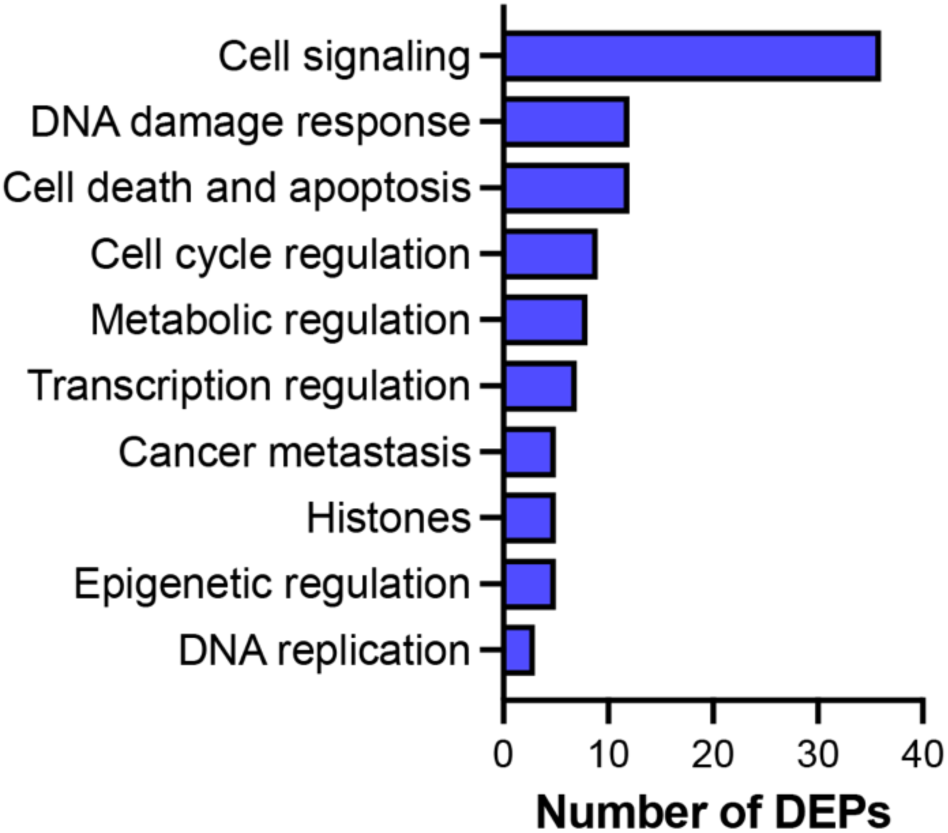
Distribution of 102 DEPs across key molecular functions. Bar graph showing the distribution of 102 DEPs categorized by their involvement in key molecular functions.

