## Supplementary Table S2 for "Therapeutic Potential of PRMT1 as a Critical Survival Dependency Target in Multiple Myeloma"

**Supplementary Table 2- Primer sequence for qRT-PCR**

| Gene | Primer sequence |
| --- | --- |
| 18s-F | GCAATTATTCCCCATGAACG |
| 18s-R | GGCCTCACTAAACCATCCAA |
| CDC45-F | GTGGGCCATCGTTGGACTAAC |
| CDC45-R | TCAAAGGAGATCCGTGTGCAG |
| PRIM1-F | ACATTCGCTACCAATCCTTCAAC |
| PRIM1-R | AGCTCCCAGCTTCACTGTATT |
| CLSPN-F | TGGAGAGTGGGGTCCATTCAT |
| CLSPN-R | CCGGGGTTTACGTTTGAAGAAA |
| TOPBP1-F | TGTGACCCTTTTAGTGGCGTT |
| TOPBP1-R | CTCTTGGGACACATCGCTGG |
| BRCA1-F | GAAACCGTGCCAAAAGACTTC |
| BRCA1-R | CCAAGGTTAGAGAGTTGGACAC |
| BRCA2-F | TGCCTGAAAACCAGATGACTATC |
| BRCA2-R | AGGCCAGCAAACTTCCGTTTA |
| FANCD2-F | ACATACCTCGACTCATTGTCAGT |
| FANCD2-R | TCGGAGGCTTGAAAGGACATC |
| FEN1-F | CACCTGATGGGCATGTTCTAC |
| FEN1-R | CTCGCCTGACTTGAGCTGT |
| ERCC4-F | TCCCTCGCCGTGTAACAAATG |
| ERCC4-R | GAGGCGCAAGATGAATGCTTC |
| RAD51-F | CAACCCATTTCACGGTTAGAGC |
| RAD51-R | TTCTTTGGCGCATAGGCAACA |
